# Inducible colistin resistance via a disrupted plasmid-borne *mcr-1* gene in a 2008 Vietnamese *Shigella sonnei* isolate

**DOI:** 10.1101/039925

**Authors:** Duy Pham Thanh, Ha Thanh Tuyen, To Nguen Thi Nguyen, Hao Chung The, Ryan R. Wick, Guy Thwaites, Stephen Baker, Kathryn E. Holt

## Abstract

The *mcr-1* gene, which confers resistance against the last-resort antimicrobial colistin, was recently discovered in Enterobacteriaceae circulating in China. Through genome sequencing we identified a plasmid-associated inactive form of *mcr-1* in a 2008 Vietnamese isolate of *Shigella sonnei*. The plasmid was conjugated into *E. coli* and *mcr-1* was activated upon exposure to colistin, suggesting the gene has been circulating in human-restricted pathogens for some time but carries a selective fitness cost.

## Introduction

The *mcr-1* gene, which confers resistance to colistin, was recently described in Enterobacteriaceae in China^1^. The index isolate, *E. coli* SHP45, was isolated at a pig farm in Shanghai in July 2013 and displayed a minimum inhibitory concentration (MIC) against colistin of 8 mg/L. *Mcr-1* was associated with a transposable element located on an IncI2 plasmid, pHNSHP45. Subsequent PCR screening using primers targeting the 5’ end of *mcr-1* detected the gene in approximately 20% of *E. coli* isolated from pigs at slaughter and 15% of *E. coli* isolated from retail meat in China. The gene was also detected amongst clinical isolates of various Gram-negative bacteria cultured from inpatients in Chinese hospitals (1.4% of *E. coli* and 0.7% of *Klebsiella pneumoniae*).

By screening available isolate collections via PCR, or mining whole genome sequence (WGS) data, many research groups have now reported the presence of *mcr-1* from widespread geographical locations and sources (food, animal and humans in Southeast Asia, Europe and Africa, and in travellers returning to Europe from Southeast Asia, South America and Africa^2-4^). The gene has been detected in multiple *Salmonella* serotypes as well as numerous *E. coli* sequence types^2^, ^5-7^ and can be carried on various plasmid backbones including IncI2, IncX4, IncHI2 and IncP^2, 7, 8^. We previously published a WGS study of ≥200 Vietnamese *Shigella sonnei* isolated from children with dysenteric diarrhoea between 1995-2010^9^. Here we report the detection of an inactivated form of the *mcr-1* gene in one of these *S. sonnei* isolates, and its selective re-activation resulting in high-level transferrable colistin resistance.

## Results

The *mcr-1* gene was detected in the genome sequence of a single *S. sonnei* strain (EG430) that was isolated in 2008 from a hospitalised child with diarrhoea in Ho Chi Minh City, Vietnam. All other sequenced *S. sonnei* isolates from the same study were negative for *mcr-1.* Assembly analysis showed the *mcr-1* gene in EG430 was associated with an IncI2 plasmid backbone, however the entire plasmid sequence could not be fully resolved using the 56 bp paired end reads^9^. Upon antimicrobial susceptibility testing *S. sonnei* EG430 was found to be susceptible to colistin (MIC 0.094 mg/L) but resistant to azithromycin (MIC 24 mg/L) via an *ermB* gene. We attributed colistin susceptibility to a 22 bp duplication of bases 503-525 of the *mcr-1* open reading frame (GAACGCCACCACAGGCAGTAAA), which induces a frameshift resulting in a truncated product (193 amino acids in length, compared to the 541 amino acid product encoded in pHNSHP45) (Fig. 1). It is also possible that a single SNP upstream of *mcr-1* (-36) in pE0G430-1 may affect expression of the gene.

**Figure 1.**
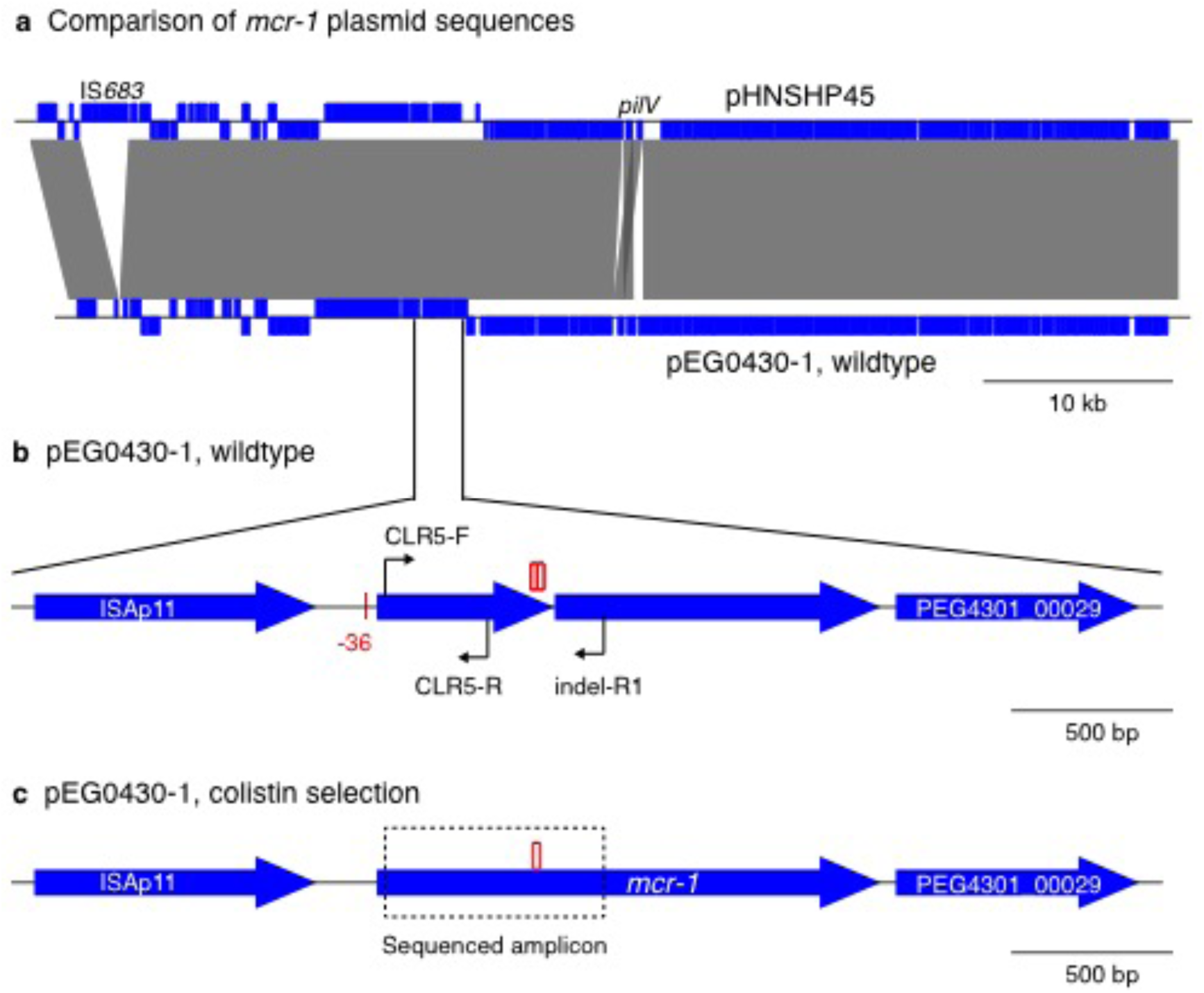
Schematic of *mcr-1* plasmid sequences. (a) Comparison of novel *mcr-1* plasmid pEG430-1 from *S. sonnei* EG430 (Vietnam, 2008) with *mcr-1* plasmid pHNSHP45 from *E. coli* (China, 2013). Blue blocks indicate protein‐coding genes; genes on the forward and reverse strands are indicated above and below the line, respectively. Grey blocks indicate regions of sequence homology between the two plasmids. (b) Zoomed in view of the *mcr-1* mobile element in pEG430-1. Large blue arrows indicate open reading frames (all are encoded on the forward strand). Arrows indicate binding sites for PCR primers. Red blocks indicate the position of 22 bp tandem repeats, present in two copies pEG430-1; red line indicates the position of a point mutation relative to pHNSHP45. (c) Mobile element with restored *mcr-1* sequence, identified in EG430 and *E. coli* transconjugants carrying pEG430-1 following selection on colistin. Dashed box shows the region that was amplified and sequenced using PCR (with primers CLR5-F and indel-R1), which confirmed restoration of the open reading frame via deletion of one copy of the 22 bp repeat.

Assuming *mcr-1* and *ermB* were located on the same plasmid we performed conjugation of *S. sonnei* EG430, using *E. coli* J53 as a recipient, and selected for transconjugants using azithromycin as a marker. PCR screening for *mcr-1* on azithromycin/sodium azide resistant *E. coli* identified multiple *mcr-1* PCR amplification-positive organisms. We subcultured the *E. coli* transconjugant, *S. sonnei* EG430 and *E. coli* J53 on LB media containing a range of colistin concentrations (0.05 mg/L to 8 mg/L). Several colonies were identified and the organisms on the highest concentration of colistin were again subcultured on media containing increasing colistin concentrations, from 8 to 32 mg/L. We were able to isolate both *E. coli* transconjugants and *S. sonnei* EG430 variants that were resistant to colistin (MIC 16 and 32 mg/L, respectively); no colonies were recovered from *E. coli* J53 at colistin concentrations above 4 mg/L. PCR amplification and sequencing of *mcr-1* using a custom primer (previously published *mcr-1* primers amplify upstream of the tandem repeat region) showed that in the colistin resistant *E. coli* and *S. sonnei* strains, one copy of the 22 bp tandem repeat had been deleted, restoring the open reading frame of *mcr-1.* We repeated this experiment multiple times and found that the re-activation of *mcr-1* was consistently reproducible.

To investigate the genetic context of *mcr-1* in detail, we isolated plasmid DNA from an *E. coli* transconjugant and sequenced it on an Illumina MiSeq to generate 250 bp paired end reads. Combined assembly of the two read sets yielded two circular plasmid sequences – pEG430-1 carrying *mcr-1,* and pEG430-2 carrying *ermB* – and not just one plasmid with both determinants as previously assumed. The pEG430-2 *ermB* encoding plasmid sequence was 68,999 bp in size, carried an IncFII *repA* gene and shared close homology with *E. coli* plasmid pHK17a (accession: JF779678.1). The pEG430-1 *mcr-1* encoding plasmid sequence was 61,826 bp in size and nearly identical to the previously described *mcr-1* encoding plasmid pHNSHP45, differing by (i) the lack of a 2,704 bp insertion of IS*683* downstream of the *repA* gene, (ii) a reorganisation of the *pilV* shufflon, and (iii) four single base substitutions, including one 36 bp upstream of *mcr-1* and (iv) the 22 bp tandem repeat in *mcr-1* (Fig. 1). IncI2 plasmid sequences were detected in 19 of the other Vietnamese *S. sonnei* isolates, however these all lacked *mcr-1* and displayed 0.3% to 1.3% nucleotide divergence from pEG430-1.

## Discussion

This is the first report describing *mcr-1* in *Shigella,* and the earliest example yet of *mcr-1* in a human clinical isolate. The only earlier example of *mcr-1* reported to date was in *E. coli* isolated from calves in France in 2005^6^, however the genetic context is not known. *E. coli* carrying *mcr-1* were recently reported in Vietnam, isolated from a pig farm in Hanoi in 2014, six years after the isolation of EG430^10^. These isolates carried *mcr-1* in distinct, non-IncI2, plasmid backbones. However, our data show the acquisition of *mcr-1* into an IncI2 plasmid backbone, and its presence in the human population in Vietnam, occurred at least as early as 2008.

Notably, the interruption we detected occurs downstream of the PCR primers used by Liu and others to detect *mcr-1^1^,* so it is possible that some isolates that test positive by PCR but not confirmed to be phenotypically resistant to colistin may carry this inactivated form of the gene. However, as we have shown, the gene can be restored again upon colistin exposure, so its presence even in inactive form could be problematic in settings where colistin is heavily used.

The fact that the *mcr-1* gene was disrupted in *S. sonnei* EG430 is concerning. *S. sonnei* is a human-restricted pathogen with no known animal reservoir, and is not likely to be regularly exposed to colistin. While the activity of Mcr-1 is not yet well understood, it appears to be a membrane-anchored enzyme with phosphoethanolamine transferase activity that likely confers resistance to colistin by a modifying lipid A. We hypothesise that in the absence of colistin exposure, these modifications carry a fitness cost and impair interactions with the human host, such that the gene may be under negative selection. The MIC we observed in *S. sonnei* carrying *mcr-1* (32 mg/L) is substantially higher than that reported previously for *mcr‐ 1* positive wildtype and transconjugant strains, which display a wide range (0.5 to 8 mg/L)^1^. We speculate that this variability in protection against colistin might be associated with the diversity of lipid A structures found in Enterobacteriaceae.

In conclusion, we have identified a deactivated version of the colistin resistance gene *mcr-1* in the human-restricted pathogen *S. sonnei,* which was isolated from a Vietnamese child in 2008. Our data suggests this gene has likely been circulating in the human population in Asia in an inducible form, suggesting a fitness cost for the active *mcr-1* gene.

## Transparency declaration

None to declare.

## Methods

### Screening for mcr-1 and IncI2 plasmids

Raw WGS data (56 bp paired-end Illumina HiSeq reads) generated previously from genomic DNA extracted from Vietnamese *S. sonnei*^9^ were screened for *mcr-1* using SRST2, which allows the detection of genes of interest direct from short reads and with higher sensitivity than assembly-based approaches^11^. IncI2 plasmids were detected by mapping reads to the pEG430-1 plasmid sequence using Bowtie2^12^. Nucleotide divergence was assessed by SNP calling using SAMtools as previously described^9^.

### Plasmid DNA extraction and sequencing

Plasmid DNA was extracted from the *S. sonnei* EG430 using the Qiagen Plasmid Midi kit (Quiagen, Germany) and sequenced via Illumina MiSeq (Illumina, USA) to generate 250 bp paired-end reads, following the manufacturer’s recommendations. Assemblies were generated using SPAdes v3.6.2^13^ and the resulting plasmid sequences were annotated using Prokka v1.11^14^. ACT (Artemis Comparison Tool)^15^ was used to compare the pEG430-1 sequence to that of the reference plasmid pHNSHP45 (accession: KP347127)^1^ and to perform manual curation of the plasmid annotations. Annotated plasmid sequences were deposited in GenBank under accessions: [TBA; these have been submitted Feb 1].

Bacterial conjugation was performed by combining equal volumes (5 ml) of overnight Luria‐Bertani (LB) cultures (approximately 5 x 10^8^ CFU/mL) of *S. sonnei* EG430 and *E. coli* J53 (sodium azide resistant). Bacteria were conjugated for 12 hours in LB broth at 37°C and *E. coli* transconjugants were selected on media containing sodium azide (100 mg/L) and azithromycin (24 mg/L). Azithromycin and sodium azide resistant *E. coli* were subjected to *mcr-1* PCR amplification to identify *mcr-1* positive transconjugants. Etests (AB Biodisk, Sweden) were used to determine the MIC of *S. sonnei* EG430, *S. sonnei* EG430 derivatives, *E. coli* J53 and E. coli J53 transconjugants against colistin. Susceptibility against colistin was determined using CLSI breakpoints^16^ (susceptible, ≤2 mg/L; intermediate, 4 mg/L; resistant ≥8 mg/L).

### PCR and sequencing

PCRs were performed using HotStar Taq DNA polymerase (Qiagen, Germany) with the recommended concentrations of reagents under the following conditions: 95°C for 15 minute, 30 cycles of 95°C for 30 seconds, 55°C for 30 seconds, 72°C for 1 minute and 72°C for 5 minutes. PCR amplification of *mcr-1* to identify transconjugants was performed using previously published primers^1^. PCR amplification of *mcr-1* to confirm copy number of the 22 bp tandem repeat in colistin resistant strains was performed using the published forward primer CLR5-F^1^ (5’-CGGTCAGTCCGTTTGTTC-3’) and a custom reverse primer MCR‐ indel-R1 (5’-TGGCTTACGCATATCAGG-3’); amplicons were sequenced using the amplification primers, using big dye terminators in both directions on an ABI 3130 sequencing machine (ABI, USA).

